# Common food preservatives impose distinct selective pressures on *Salmonella* Typhimurium planktonic and biofilm populations

**DOI:** 10.1101/2023.09.28.559892

**Authors:** Justin Abi Assaf, Emma R. Holden, Eleftheria Trampari, Mark A. Webber

**Author notes:** Corresponding authors: Eleftheria Trampari, Mark Webber. These authors contributed equally to this work (Justin Abi Assaf, Emma R. Holden). **Competing interests** The authors declare no competing interests.

## Abstract

Food preservatives are crucial in controlling microbial growth in processed foods to maintain food safety. Bacterial biofilms pose a threat in the food chain by facilitating persistence on a range of surfaces and food products. Cells in a biofilm are often highly tolerant of antimicrobials and can evolve in response to antimicrobial exposure. Little is known about the efficacy of preservatives against biofilms and their potential impact on the evolution of antimicrobial resistance. In this study we investigated how the common food pathogen *Salmonella enterica* serovar Typhimurium responded to subinhibitory concentrations of four common food preservatives (sodium chloride, potassium chloride, sodium nitrite or sodium lactate) when grown planktonically and in biofilms. We found that each preservative exerted a unique selective pressure on *S*. Typhimurium populations grown planktonically and in a biofilm. Biofilm formation itself seemed to confer protection when exposed to each of the four preservatives, more so than previous exposure to sub-inhibitory concentrations of preservatives. There was a trade-off between biofilm formation and growth in the presence of three of the four preservatives, where prolonged preservative exposure resulted in reduced biofilm biomass and matrix production over time. Despite the differences in biofilm formation and preservative tolerance seen following three preservative stresses, they selected for mutations in global stress response regulators *rpoS* and *crp*. There was no evidence for any selection of cross-resistance to antibiotics after preservative exposure, and some evidence that antagonism between preservatives can be exploited in compound cocktails to reduce contamination in the food chain.

**Highlights:**

- Preservative-specific evolutionary adaptation of *Salmonella* was shown over time.
- A trade-off between adaptation and biofilm formation was observed.
- No cross-resistance to antibiotics was seen after preservative exposure.
- Mutations were found to be preservative-specific, with some common ones like *rpoS* and *crp*.

## Introduction

Food preservatives are routinely used to protect processed food from microbial contamination and their effectiveness against spoilage has been repeatedly demonstrated. There are several preservatives used in the food chain, the efficacy and specificity of which is dependent on several factors including temperature, pH, and initial bacterial load (Brul and Coote, 1999). Antimicrobial preservatives such as sodium chloride (NaCl) and potassium chloride (KCl) act by reducing water availability in food limiting microbial growth whilst also imposing osmotic stress on bacteria (Li et al., 2021, Guan et al., 2017, Henney et al., 2010). Sodium lactate (SL) inhibits bacterial growth through creating low pH and sodium nitrite (SN) inhibits growth due to generation of peroxynitride which is a cytotoxic biological agent causing the oxidation and nitration of cell components including DNA, proteins and lipids (Majou and Christieans, 2018). These preservatives are in common use due to their cost effectiveness and activity against several pathogenic bacteria including *Salmonella, Escherichia coli* and *Clostridium* species.

Despite the frequent use of chemical preservatives in the food industry, relatively little research has studied their mechanistic effects on bacterial growth and examined the potential of evolution of bacterial resistance.

*Salmonella enterica* is a leading foodborne pathogen having a significant socio-economic cost. Although in recent years, there is a significant decrease in the use of antimicrobials in agriculture, emergence of antimicrobial resistance is an ongoing threat. Approximately 40% of European *Salmonella enterica* serovar Typhimurium isolates are now found to be resistant to multiple antibiotics (EFSA and ECDC, 2018). It has also been shown that non-antibiotic antimicrobials can select for cross-resistance to antibiotics (Webber et al., 2015) making the use of antimicrobials in a non-clinical context important to understand.

In our recent work, we developed a biofilm evolution model to study how *S*. Typhimurium biofilms evolve in response to antimicrobial stress (Trampari et al., 2021). Bacteria are commonly found in biofilms, aggregated communities of cells encased in a matrix that allows their attachment to surfaces (Berlanga and Guerrero, 2016, Darby et al., 2023). Bacteria within a biofilm can often tolerate large changes in pH, osmolarity, lack of nutrients, external mechanical forces, and higher concentrations of antibiotics relative to their planktonic counterparts. This resistance to stress is mediated by the biofilm’s extracellular matrix and the changes in physiological state seen for cells within a biofilm (Singh et al., 2021). Biofilms pose a challenge to the food industry as they allow bacteria to bind to a range of surfaces and food products, making decontamination more difficult (Carrascosa et al., 2021). We previously showed that biofilms can rapidly develop antibiotic resistance but that this comes at a cost to fitness, virulence, and biofilm formation itself (Trampari et al., 2021). In this work, we used this biofilm evolution model to investigate the genotypic and phenotypic effects of continuous exposure to subinhibitory concentrations of common food preservatives; sodium chloride, potassium chloride, sodium lactate and sodium nitrite, on the foodborne pathogen *Salmonella enterica* serovar Typhimurium. We found that these preservatives affected biofilm formation and growth rate in a compound-specific manner. The ability of *S*. Typhimurium to form biofilms was significantly reduced following continuous exposure to NaCl, KCl and SL, but was unaffected by SN. We identified emergence of cross-preservative tolerance between populations exposed to NaCl and SN and an antagonistic effect of KCl and SL on *S*. Typhimurium growth. There was no cross resistance observed against antibiotics after exposure to any of the preservatives. Whole genome sequencing identified selection of mutations which were largely preservative-specific although mutations in stress response regulators *crp* and *rpoS* were shared between two or more preservatives. Deletion of these genes confirmed roles influencing fitness under preservative stress.

## Methods

### Biofilm evolution model

Each preservative stress condition was made up of eight lineages: two preservative-exposed planktonic lineages (P1 and P2), two unexposed biofilm lineages (C1 and C2), and four exposed biofilm lineages (A, B, C, D) (Trampari et al., 2021). Biofilm lineages were comprised of three 5 mm stainless steel beads (from Simply Bearings) in 5 mL LB broth without salt. Planktonic lineages were set up the same but did not contain beads. Approximately 10^5^ CFU/mL of *Salmonella enterica* subsp. *enterica* serovar Typhimurium strain 14028*S* was inoculated into each condition and cultures were incubated at 30°C for 72 hrs horizontally with mild agitation at 40 rpm. Cultures were exposed to preservatives at subinhibitory concentrations, corresponding to 5% NaCl, 7 % KCl, 5 % SL or 0.06 % SN. Subinhibitory concentrations were selected as those that inhibited the growth of *S*. Typhimurium by 25-35 %.

Cultures were passaged into new media every 72 hours. For biofilm lineages, one steel bead was washed in sterile PBS and transferred to fresh media to seed the next passage. For planktonic conditions, 5 μL of culture was added to fresh media. A sample of each condition was stored at each passage: this consisted of 1 mL of planktonic culture or one bead from the biofilm lineages. Beads were washed with sterile PBS to remove planktonic cells and vortexed in fresh sterile PBS to resuspend the biofilm. Both planktonic cultures and resuspended biofilms were mixed with 7.5 % DMSO and stored at -20 °C. Biofilm generation number was calculated by multiplying the passage number by log_2_ of the dilution factor (average number of cells recovered per bead for biofilms).

### Growth kinetics

Populations were grown in the presence of preservatives to investigate preservative tolerance and cross-tolerance. Approximately 10^5^ CFU/mL of liquid culture was added to LB broth in a 96-well plate supplemented with either 5 % NaCl, 7 % KCl, 5 % SL or 0.06 % SN. A wild type *S*. Typhimurium 14028*S* was included on each plate as a control. Cultures were grown for 20 hours at 37 °C in a FLUOstar Omega plate reader (BMG Labtech) and optical density of the culture (OD 600 nm) was measured every 15 minutes. The area under each growth curve was calculated in R (version 4.2.1) using the AUC function in the DescTools package.

### Measuring biofilm biomass and matrix composition

Biofilm formation was quantified by measuring biofilm biomass and production of curli, an amyloid protein in the biofilm matrix. To quantify biofilm biomass, approximately 10^5^ CFU/mL of liquid culture was added to a polystyrene 96-well plate of LB broth without salt and incubated at 30 °C for 48 hours. Planktonic cells were rinsed away with water and 0.1 % crystal violet (w/v) was added to the plate for 10 minutes to stain the biofilm. Plates were rinsed with water to remove unbound dye, and the stained biofilm was solubilised with 70 % ethanol. Biofilm biomass was quantified using a FLUOstar Omega plate reader (BMG Labtech) at an absorbance wavelength of 590 nm. Data shown consists of four independent replicates, and a wild type *S*. Typhimurium 14028*S* was included on each plate as a control. Curli production was measured by spotting 5 μL of approximately 10^7^ CFU/mL of liquid culture on LB agar without salt containing 40 μg/mL Congo red. Congo red binds to curli, an amyloid protein in the *S*. Typhimurium biofilm matrix. Plates were photographed and colonies shown are representative of a minimum of four independent replicates.

### Antimicrobial susceptibility testing

A standard two-fold agar dilution method was used to determine the MICs of different antibiotic classes against the various lineages to find out whether preservative exposure could lead to a cross-resistance to antimicrobials. Mueller-Hinton agar was supplemented with chloramphenicol, tetracycline, azithromycin, ciprofloxacin, kanamycin, cefotaxime, ampicillin or colistin. Bacterial cultures were diluted to approximately 10^7^ CFU/mL and spotted onto agar plates using a 96-pin replicator. Plates were incubated overnight at 37 °C and the minimum inhibitory concentration (MIC) was determined as the lowest concentration of drug tested that inhibited or stopped the bacterial growth. Data shown represents two biological and two technical replicates, and a wild type was included on each plate as a control.

### DNA sequencing and SNP analysis

DNA was isolated from planktonic and biofilm cultures and sequenced following the protocol described by Trampari et al. (2021). SNPs were identified by comparing FASTQ files from each isolate to *S*. Typhimurium 14028*S* reference genome CP001363 (Jarvik et al., 2010) using Snippy version 4.6 (https://github.com/tseemann/snippy).

### Gene deletion mutant creation

Single gene deletion mutants were created in *S*. Typhimurium following the gene doctoring protocol (Lee et al., 2009) using plasmids created using Golden gate assembly (Thomson et al., 2020). Gene deletions were validated via Sanger sequencing of the region of interest as well as Illumina whole genome sequencing as described above.

## Results

### Preservative exposure selects for changes in growth rate in a compound-specific manner

Populations of *S*. Typhimurium were grown as biofilms on stainless steel beads. Four independent lineages (denoted A, B, C and D) were exposed to either 5% NaCl, 7% KCl, 5% SL or 0.06% SN. These concentrations were selected as they caused a 25-35% reduction in planktonic growth rates representing a stress which will allow a population to reach stationary phase but impose a selective pressure. Two independent unexposed biofilm controls and two exposed planktonic controls were also included. Cultures were passaged every 72 hours into fresh media: for planktonic cultures this used a 1:10,000 dilution, and for biofilms, one stainless steel bead was transferred to fresh LB broth containing sterile beads to seed the next generation. Populations were repeatedly exposed until improvements to growth rates plateaued. For NaCl and KCl this was after 48 passages (representing approximately 1000 generations for biofilms) and for sodium lactate and sodium nitrite this was 24 generations (approximately 500 generations in biofilms).

Cells were recovered from the biofilm populations at each passage and tested for their ability to grow in the presence of the preservative they were exposed to (Figure 1). Exposure to NaCl for approximately 500 generations resulted in reduced growth in all populations at the middle time point relative to the early time point in the presence of NaCl (*p* < 0.0001) (Figure 1a). Following 1000 generations of exposure to NaCl, biofilm populations grew significantly worse in the presence of NaCl at the late time point relative to unexposed biofilm controls (*p* = 0.006) and relative to preservative-exposed planktonic cultures (*p* < 0.0001). This suggests a trade-off between biofilm formation and growing in the presence of the preservative, whereby individually they confer reduced susceptibility to the preservative, but populations cannot adapt under both conditions simultaneously.

**Figure 1:**
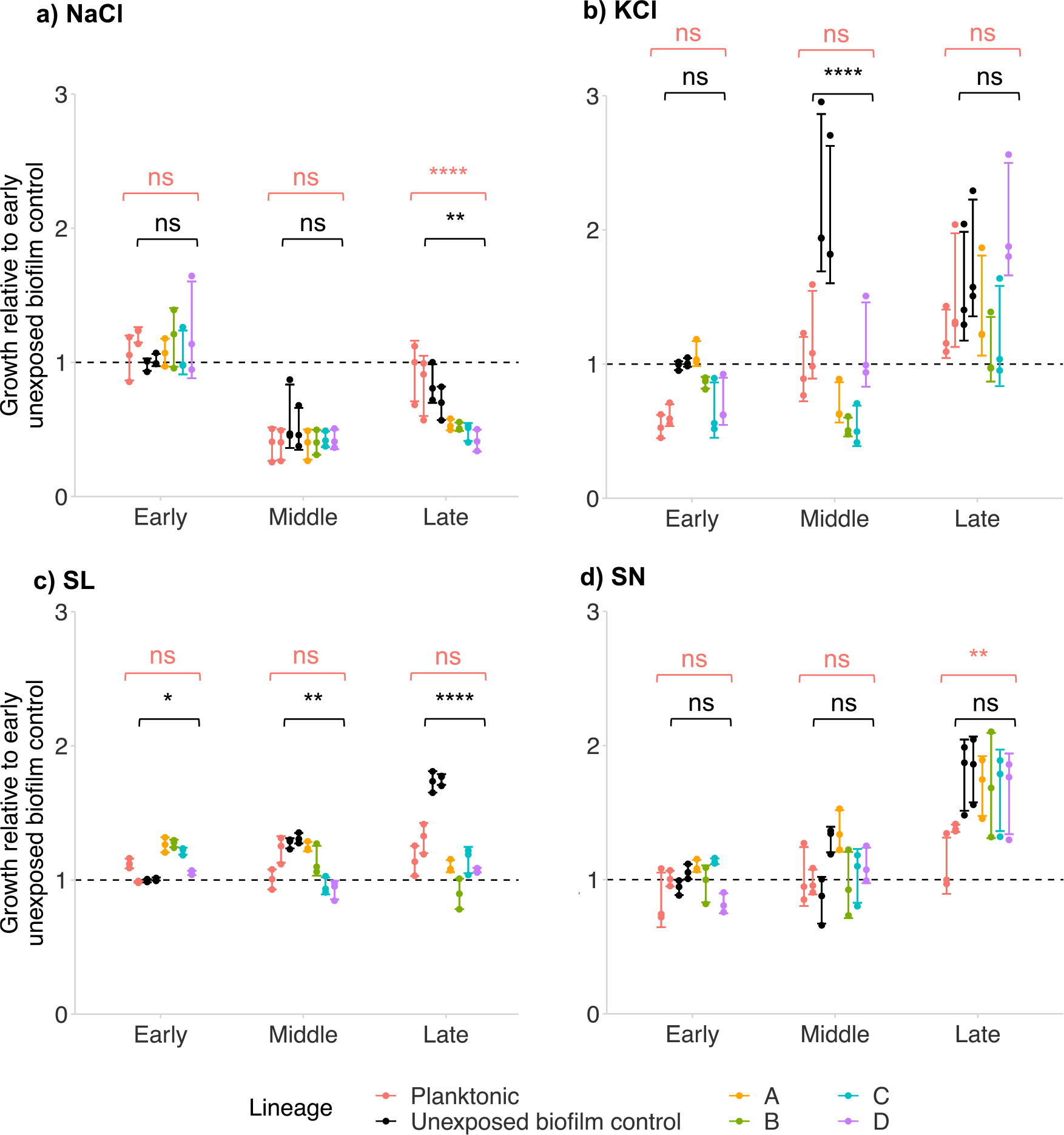
Growth of populations continuously exposed to **a)** sodium chloride (NaCl), **b)** potassium chloride (KCl), **c)** sodium lactate (SL) and **d)** sodium nitrite (SN), grown in the presence of the preservative they were exposed to, relative to unexposed controls at the early time point. Four independent biofilm lineages (A, B, C and D, separated by colour), two unexposed biofilm control linages and two preservative-exposed planktonic lineages were exposed for 24 or 48 passages with isolates taken at ‘early’, ‘middle’ and ‘late’ stages (corresponding to passage numbers 1, 25, 48 for NaCl and KCl; and passage numbers 1, 12, 24 for SN and SL). Growth represents the area under growth curves created by measuring the optical density (Absorbance 600 nm) of cultures over 18 hours. Points show three independent replicates and error bars denote one standard deviation. Asterisks show significant differences (Two-way ANOVA with Tukey post hoc analysis) between the preservative-exposed biofilm lineages and the unexposed biofilm controls (in black) or the preservative-exposed planktonic lineages (in red): ns not significant; * *p* < 0.05; ** *p* < 0.01; *** *p* < 0.001; **** *p* < 0.0001.

Growth in the presence of KCl increased over time for all populations: there was a significant increase in growth from the early to the late time points for planktonic (*p* = 0.003), unexposed biofilm (*p* < 0.02) and preservative-exposed biofilm populations (*p* = 0.0009) (Figure 1b). This suggests that preservative exposure and biofilm formation both separately and together confer protection against preservative killing. At the middle time point (approximately 500 generations), growth of unexposed biofilms in the presence of KCl was significantly higher than biofilms continuously exposed to KCl (*p* < 0.0001). After 1000 generations, there was no significant difference in growth between unexposed biofilm populations and preservative-exposed biofilm and planktonic populations in the presence of KCl.

Biofilm populations exposed to SL had increased growth in the presence of SL relative to unexposed biofilms at the early time point, following one passage in SL (*p* = 0.017) (Figure 1c). However over evolutionary time, growth of exposed biofilms in the presence of SL remained relative unchanged whereas growth of unexposed biofilms increased and was significantly greater than exposed biofilms at the middle (*p* = 0.005) and late (*p* < 0.0001) time points. It appears that prolonged exposure to SL does not affect the ability to either planktonic or biofilm populations to grow in the presence of SL, but biofilm formation without preservative stress reduces susceptibility to SL. Similar to the relationship seen with NaCl, this suggests a trade-off between biofilm formation and preservative adaptation, where biofilm formation confers reduced susceptibility to SL but prolonged preservative exposure prevents this adaptation in biofilm populations.

SN-exposed and unexposed biofilm populations had increased growth in the presence of SN following 500 generations of exposure at the late time point relative to the early time point (*p* < 0.0001), whereas the growth of planktonic populations remained relatively unchanged over evolutionary time (Figure 1d). Similar to the relationship seen with SL, exposure to SN does not seem to affect the ability of populations to grow in the presence of SN and biofilm formation appears to confer more protection in the presence of SN, irrespective of whether that biofilm population has previously been exposed to it.

### Continuous exposure to preservatives affects biofilm formation in *S*. Typhimurium in a compound- specific manner

We investigated how exposure to different preservatives over time affected the ability of *S*. Typhimurium to form a biofilm. Biofilms were grown for 48 hours on polystyrene 96-well plates and stained with crystal violet to quantify biomass relative to wild type *S*. Typhimurium. Cultures were also spotted on agar containing Congo red, which binds to an amyloid protein in the biofilm matrix called curli, allowing a visual marker of curli production in the biofilm. Biofilm characteristics were measured at the ‘early’, ‘middle’, and ‘late’ time points relative to generation time in the population (corresponding to passage numbers 1, 25, 48 for NaCl, KCl and unexposed controls; and passage numbers 1, 12, 24 for SN and SL).

Biofilm biomass increased relative to the wild type over time in the unexposed biofilm populations, showing the model itself selects for increased biomass production in the absence of any stressor (figure 2a). Populations continuously exposed to NaCl, KCl or SL showed a significant reduction in biofilm biomass over time relative to early time point (figure 2 b,c,d). This reduction in biofilm biomass was paired with a reduction in curli production in the biofilm and suggests continuous exposure to these preservatives reduces the ability of *S*. Typhimurium to form biofilms. However, this was not the case following exposure to SN, which had no significant effect on biofilm biomass or curli production over time (figure 2e). Whilst biomass was reduced following exposure to NaCl, KCl or SL, the productivity (e.g. number of cells produced by a biofilm) of biofilm populations exposed to each preservative was unchanged from unexposed biofilm cultures (figure 2f), suggesting preservative exposure changes the nature of the biofilms formed and results in less elaborate matrix production but does not significantly reduce the biofilm population size produced.

**Figure 2:**
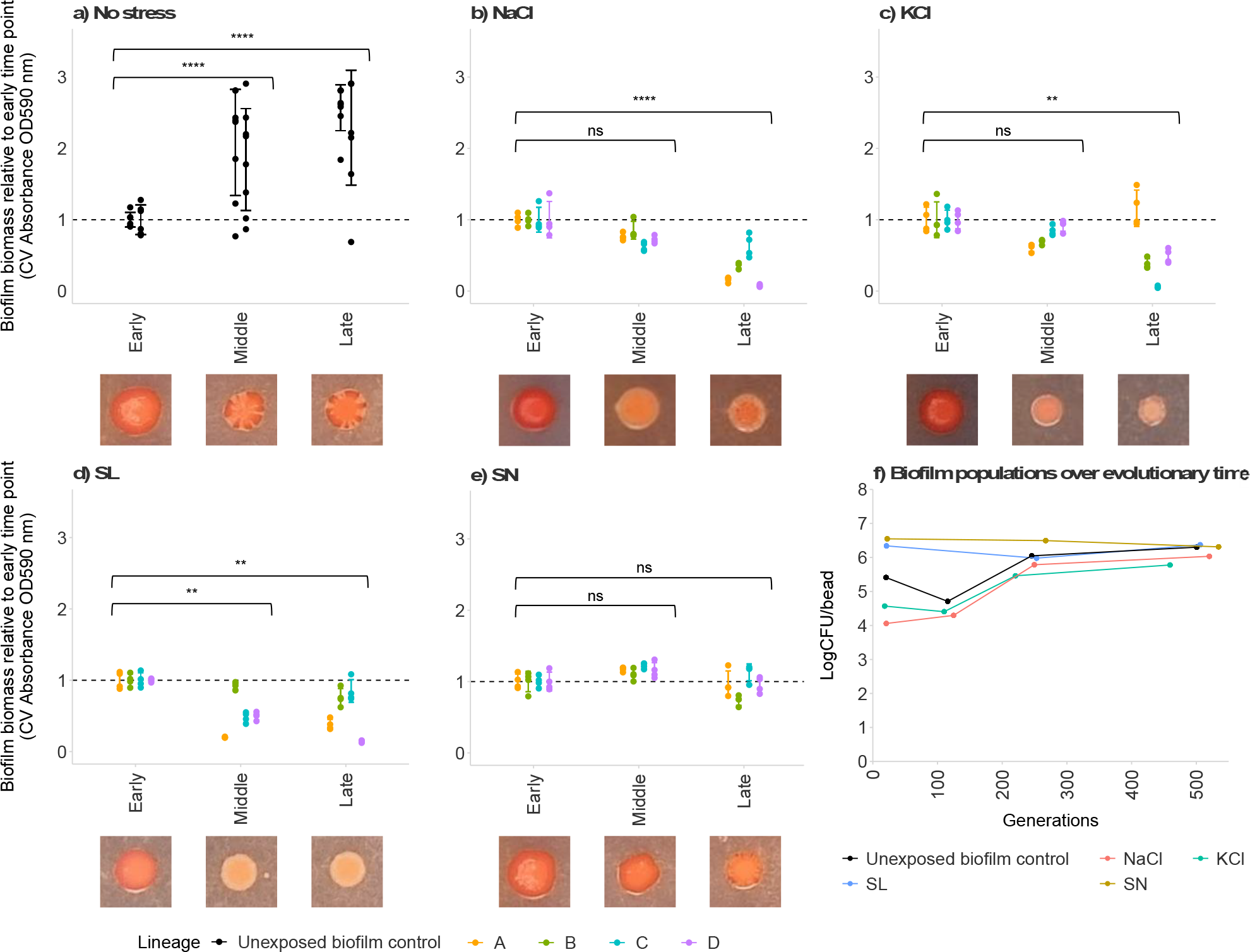
Biofilm formation of *S*. Typhimurium continuously exposed to **a)** no stress, **b)** sodium chloride (NaCl), **c)** potassium chloride (KCl), **d)** sodium lactate (SL) and **e)** sodium nitrite (SN). Four independent lineages (A, B, C and D, separated by colour) and two unexposed control lineages were stressed for 24 or 48 passages with isolates taken at ‘early’, ‘middle’ and ‘late’ stages (corresponding to passage numbers 1, 25, 48 for NaCl, KCl and unexposed controls; and passage numbers 1, 12, 24 for SL and SN). Biofilm biomass was measured by crystal violet (CV) staining and is shown relative to the early time point. Points show relative biofilm biomass of four independent replicates and error bars represent one standard deviation. Statistical significance (Two-way ANOVA with Tukey post hoc analysis) between biofilm populations at each time point is denoted by asterisks: * *p* < 0.05, ** *p* < 0.01, *** *p* < 0.001, **** *p* < 0.0001, ns not significant. Curli in the biofilm matrix was visualised by staining with Congo red. Images are representative of two independent replicates. **F)** Biofilm density on steel beads over evolutionary time as populations of S. Typhimurium were continuously exposed to NaCl, KCl, SN and SL. The number of generations at each time point was calculated using the number of passages and the average number of cells recovered per bead for that condition. Points and lines show the average biofilm density of four technical replicates from each of the four populations exposed to each preservative, and the shaded ribbon denotes 1 standard deviation from the mean.

### Preservatives do not select for cross-tolerance to antibiotics in S. Typhimurium biofilms

Populations continuously exposed to each of the four food preservatives were tested for any change in antibiotic susceptibility. There were no significant changes in susceptibility to chloramphenicol, tetracycline, azithromycin, ciprofloxacin, kanamycin, cefotaxime, colistin or ampicillin in any of the populations tested (supplementary table 1). This indicates that continuous exposure to these four preservatives does not confer reduced susceptibility to antibiotics.

### Preservatives select for cross-tolerance to other preservatives

We also investigated preservative cross-tolerance by growing biofilm populations continuously exposed to one preservative in the presence of the others for 20 hours. Biofilm populations continuously exposed to NaCl had a significantly increased growth relative to the wild type when grown in the presence of NaCl (*p* = 0.020) or SN (*p* < 0.0001), but not KCl or SL (figure 3a). This may suggest a shared mechanism affecting preservative susceptibility in biofilms exposed to NaCl or SN. Continuous exposure to KCl significantly reduced growth in the presence of KCl relative to the wild type (*p* = 0.015), but growth was unchanged from the wild type in the presence of NaCl, SL or SN (figure 3b). It appears prolonged exposure to KCl does not affect the ability of biofilm populations to grow in the presence of any of the other preservatives tested. There was significantly reduced growth of biofilm populations continuously exposed to SL when grown in the presence of NaCl (*p* = 0.033), SL (*p* = 0.003) or SN (*p* < 0.0001), but not KCl (figure 3c). This may indicate prolonged exposure to SL affects growth in the presence of preservatives containing sodium. Biofilm populations continuously exposed to SN had no significant difference in growth relative to the wild type in the presence of any of the four preservatives tested (figure 3d).

**Figure 3:**
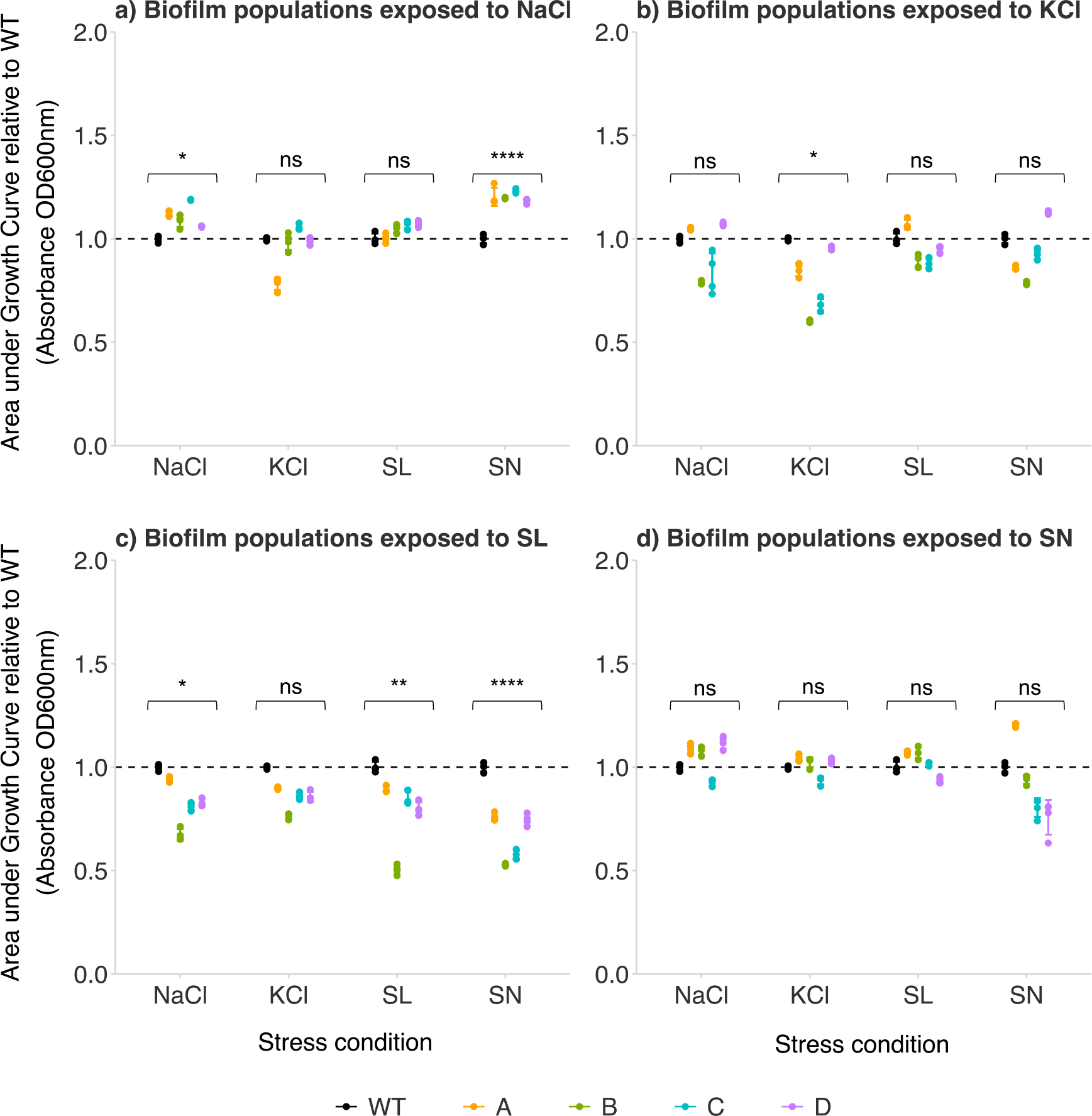
Growth of late-stage biofilm populations continuously exposed to **a)** sodium chloride (NaCl), **b)** potassium chloride (KCl), **c)** sodium lactate (SL) and **d)** sodium nitrite (SN) relative to wild type (WT) S. Typhimurium when grown in the presence of each of the four preservatives for 20 hours. Four independent lineages (A, B, C and D, separated by colour) were continuously exposed to NaCl or KCl for 48 passages (1000 generations), or SL or SN for 24 passages (500 generations). Growth rates were determined by calculating the area under each growth curve and normalising to the WT grown in the same condition. Points show growth of four independent replicates of each of the four independent lineages (A, B, C and D, separated by colour). Error bars denote 1 standard deviation and asterisks show statistical significance (Two-way ANOVA with Tukey post hoc analysis) between the wild type and preservative-exposed biofilm populations cultured under each stress: * *p* < 0.05, ** *p* < 0.01, *** *p* < 0.001, **** *p* < 0.0001, ns not significant.

### Whole genome sequencing of preservative exposed populations revealed mutations selected by preservative exposure

To investigate the genetic basis for the altered phenotypes observed, populations for each preservative stress in all four exposed biofilm lineages were whole genome sequenced. Unexposed biofilm populations and exposed planktonic populations were also sequenced as controls. Preservative exposure was found to select for SNPs within genes involved in fimbriae production, sugar degradation, respiration, LPS biosynthesis, cell division, transmembrane transport as well as multiple different stress response mechanisms (figure 4). There were more SNPs identified in exposed biofilm lineages relative to their planktonic counterparts (figure 4b). No common mutations were identified to be shared among the four exposures, suggesting no single mechanism for surviving preservative stress.

**Figure 4:**
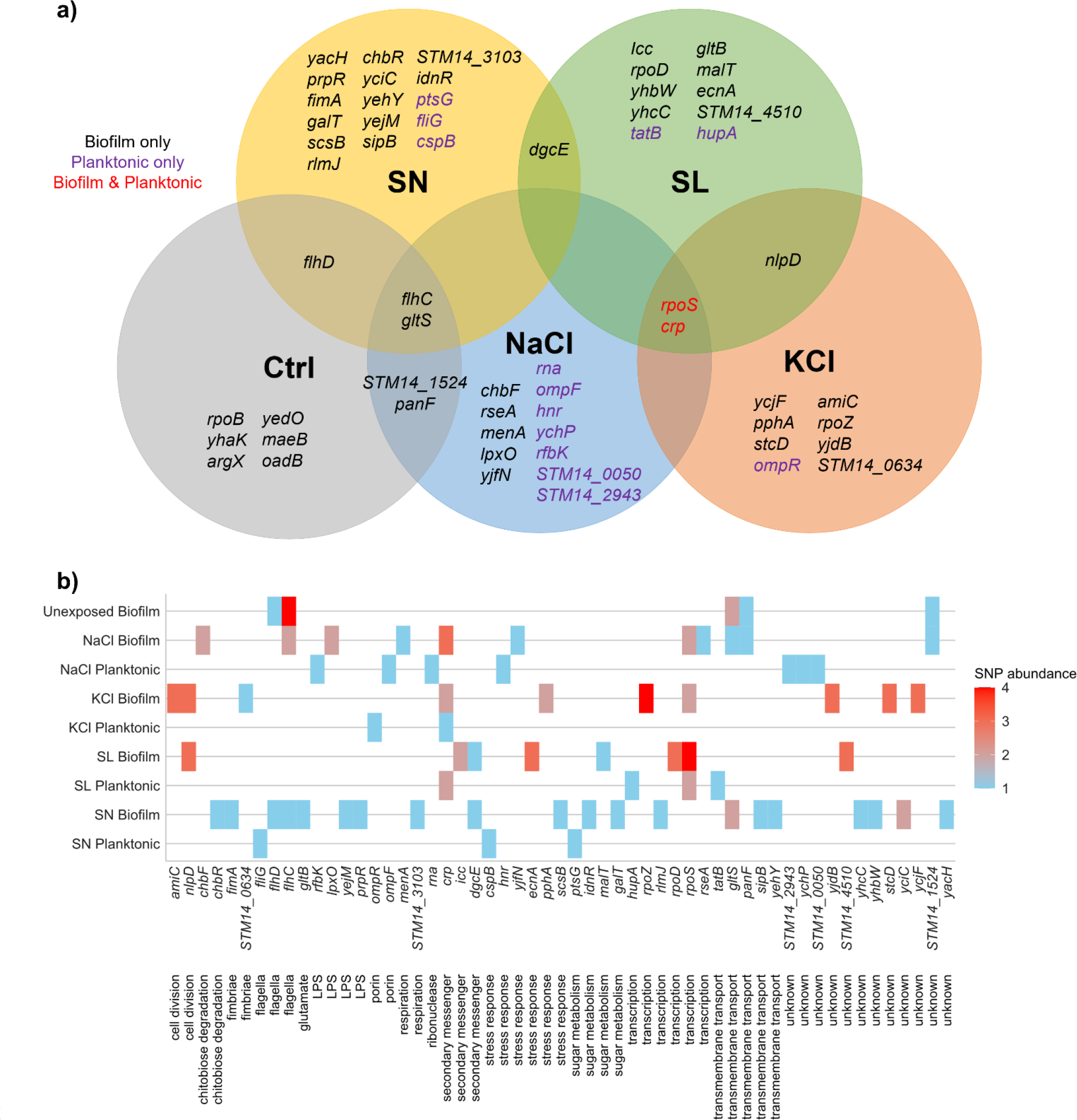
**a)** Genes in *S*. Typhimurium in which SNPs were identified following continuous exposure to sodium chloride (NaCl), potassium chloride (KCl), sodium lactate (SL) and sodium nitrite (SN) in biofilm and planktonic cultures, as well as in unexposed biofilm controls (Ctrl). SNPs found in the biofilm lineages, planktonic lineages or both have been distinguished by colour. **B)** SNP abundance across the four exposed biofilm lineages, two exposed planktonic lineages and two unexposed biofilm controls, combined for each preservative stress. Further details of the lineage and SNP position are detailed in supplementary table 2.

There were no common SNPs shared between both planktonic and biofilm lineages exposed to NaCl or SN (figure 4b), suggesting these preservatives exert a distinct selective pressure on planktonic and biofilm populations. Following continuous exposure to KCl, same missense variant of *crp* (C19Y) was seen in both biofilm and planktonic cultures, as well as in biofilms continuously exposed to NaCl and planktonic populations exposed to SL. This suggests this specific mutation within *crp* is beneficial for survival in the presence of a broad range preservatives but is not specific to biofilm populations. Cyclic AMP receptor protein (encoded by *crp*) and the secondary messenger molecule cAMP are involved in regulating respiration, stress responses and virulence, alongside *rpoS* (Donovan et al., 2013). Mutations in *rpoS* were seen in biofilms exposed to NaCl or KCl, as well as both planktonic and biofilm populations exposed to SL. This gene encodes sigma factor S responsible for regulating the cell’s general stress response (Schellhorn, 2014). Each condition and preservative exposure resulted in different SNPs in different locations within *rpoS*. As well as roles in stress responses both *crp* and *rpoS* have well defined roles in biofilm formation in *S*. Typhimurium providing a potential mechanistic link between adaptation to the preservative stress with changes in biomass production described above (Holden et al., 2022).

To explore the idea that loss of function of *crp* or *rpoS* would impact both biofilm formation and preservative tolerance, we created defined deletion mutants of each in *S*. Typhimurium. Deletion of *crp* did not significantly affect biofilm biomass in the presence or absence of any preservative stress (figure 5a). Growth of a *crp* deletion mutants was significantly reduced in the presence of KCl relative to the wild type (*p* = 0.014), but other preservatives had no relative effect on growth (figure 5b). This suggests that the SNPs within *crp* selected under preservative stress do not confer a loss of function, as deletion of *crp* does not replicate the effect on biofilm formation and preservative susceptibility seen following prolonged preservative exposure.

**Figure 5:**
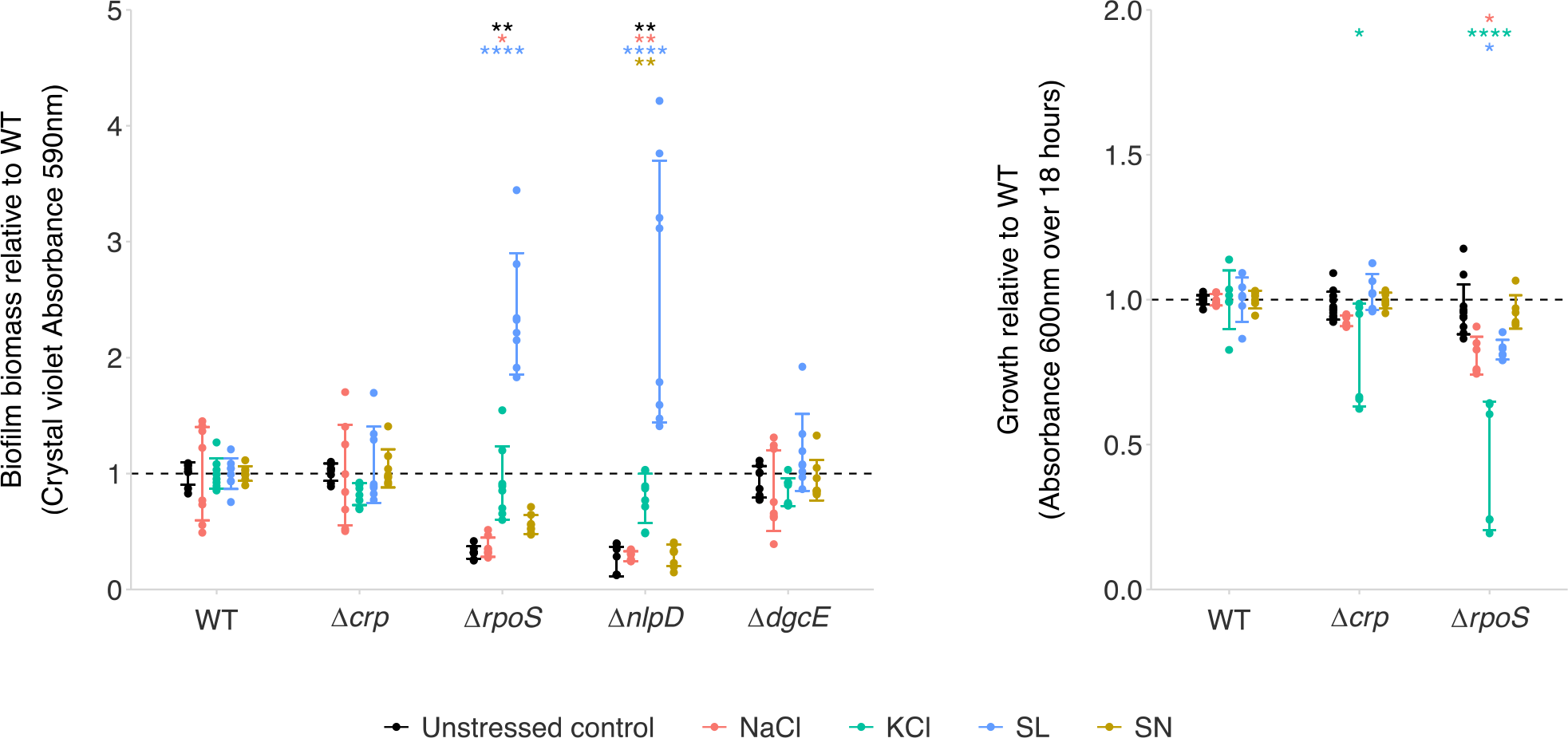
**a)** Biofilm formation of single gene deletion mutants relative to wild type (WT) *S*. Typhimurium grown in the presence of sodium chloride (NaCl), potassium chloride (KCl), sodium lactate (SL) and sodium nitrite (SN). Points show biofilm biomass measured via crystal violet staining of two biological and four technical replicates. **B)** Growth of single gene deletion mutants relative to wild type (WT) *S*. Typhimurium in the presence of the same four preservatives. Growth represents the area under growth curves created by measuring the optical density (Absorbance 600 nm) of cultures over 18 hours. Points show two biological and three technical replicates. For both panels, error bars show one standard deviation and asterisks show significant differences (Two-way ANOVA with Tukey post hoc analysis) between the wild type and the single gene deletion mutants under the same preservative stress, distinguished by colour: ns not significant; * *p* < 0.05; ** *p* < 0.01; *** *p* < 0.001; **** *p* < 0.0001.

Deletion of *rpoS* resulted in significantly reduced biofilm biomass relative to the wild type in the absence of any preservative stress (*p* = 0.009) and in the presence of NaCl (*p* = 0.023), but significantly increased biofilm biomass in the presence of SL (*p* < 0.0001) relative to the wild type (figure 5a). This suggests loss of RpoS function can be conditionally beneficial to biofilm formation, but there is no single common mechanism affecting susceptibility to all preservatives. Despite conditional effects on biofilm formation, growth of a *rpoS* deletion mutant was significantly reduced in the presence of NaCl (*p* = 0.012), KCl (*p* < 0.0001) and SL (*p* = 0.044) relative to the wild type (figure 5b).

Different SNPs were also identified in *nlpD* in biofilms continuously exposed to KCl or SL (figure 4). This gene is involved in septal splitting for cell division alongside *amiC* (13), in which a SNP was identified in biofilms exposed to KCl. This implies that cell division affects survival in the presence of KCl and SL in the biofilm. Deletion of *nlpD* resulted in reduced biofilm biomass without preservative exposure relative to the wild type (*p* = 0.001), and further reduced biomass upon exposure to NaCl (*p* = 0.004) or SN (*p* = 0.005) (figure 5a). In biofilms exposed to SL, a Δ*nlpD* mutant had increased biofilm biomass relative to the wild type (*p* < 0.0001), similar to the Δ*rpoS* mutant biofilm (figure 5a). This may indicate a shared regulator or function in response to preservative stress.

In biofilm populations continuously exposed to SN or SL, two separate missense mutations were found in *dgcE*, encoding a diguanylate cyclase involved in c-di-GMP synthesis (figure 4). C-di-GMP has a well-establish role in biofilm formation (Hengge, 2020), however mutations within *dgcE* were only found in preservative-exposed biofilms, not in unexposed biofilms, suggesting a specific role in stress response in the biofilm. However, deletion of *dgcE* resulted in no significant difference in biofilm biomass production in the presence or absence of the preservatives tested.

Various mutations were found in both unexposed and preservative-exposed biofilms: these were located in *flhC, flhD, gltS, panF* and *STM14_1524*. Mutations within these genes are likely a result of selection for continuous biofilm formation rather than a preservative-specific response. FlhDC is a master regulator of flagella biosynthesis and function with a complex role in biofilm formation. Spatial and temporal regulation of *flhDC* expression is essential for the development of the biofilm through time (Samanta et al., 2013, Holden et al., 2021). Both *gltS* and *panF* encode transmembrane symporters, specifically a glutamate:sodium symporter and a pantothenate:cation (Na^+^) symporter, respectively. Previous work has shown deletion of *gltS* in *E. coli* reduced biofilm biomass, and neither *panF* or *STM14_1524* have previously been linked to biofilm formation.

## Discussion

This work shows that four common preservatives impose distinct selective pressures on *S*. Typhimurium. Planktonic and biofilm cultures adapt to subinhibitory concentrations of KCl or SN and have increased growth in the presence of these preservatives following prolonged exposure. However, adaptation is not seen in the presence of NaCl or SL, where populations continuously exposed to these preservatives had reduced growth or unchanged growth respectively over time.

We described how preservative exposure comes at a cost to biofilm formation, whereby biofilm biomass and curli production were reduced following prolonged exposure to NaCl, KCl, or SL, but unchanged for populations exposed to SN. Previous work by Lamas et al. (2018) also found no significant change in biofilm formation by many *Salmonella* serovars on stainless steel following exposure to sodium nitrite. However, sodium nitrite was found to reduce biofilm formation in some Gram positive bacteria such as *Staphylococcus* and *Streptococcus* (Schlag et al., 2007, Al-Ahmad et al., 2008). Whilst biofilm biomass production was reduced, the productivity of biofilms formed on steel beads over time did not change. This indicates that biofilm cell numbers and productivity are unchanged, but biofilm matrix production is considerably affected.

There was some evidence of cross-tolerance between preservatives, where prolonged exposure to NaCl preservative improved survival in the presence of SN relative to the wild type. The two genes containing SNPs common to both NaCl- and SN-exposed biofilm populations were also common to unexposed biofilm populations, suggesting biofilm formation itself is the mechanisms that confers cross-tolerance to these preservatives. This is supported by our finding that unexposed biofilm populations had increased growth in the presence of NaCl relative to those that were exposed to NaCl for 1000 generations, and biofilm populations had increased growth in the presence of SN relative to planktonic culture irrespective of their SN exposure.

Prolonged exposure to SL resulted in reduced growth in the presence of NaCl, SL and SN, but not KCl. Interestingly, there were no SNPs in genes common to these three sodium-containing preservatives that were not found populations exposed to KCl. This suggests no common mechanism and that SL selects for mutations in multiple genes that affect susceptibility to a wide range of preservatives. Further investigation into the mechanisms of preservative tolerance is necessary to understand the implications and mechanisms of this relationship.

Each preservative selected a unique set of mutations in S. Typhimurium populations with only a small number shared across each preservative. Mutations within *crp* and *rpoS* were seen in populations exposed to three of the four preservative stresses in both planktonic and biofilm lineages. Loss of function mutations and insertions within *rpoS* have been found to increase survival in the presence of trimethoprim (Turner et al., 2021). Additionally, mutations in *crp* and another gene involved in cAMP activity (*cyaA*) were found to reduce susceptibility to 22 antibiotics (Alper and Ames, 1978) and quaternary ammonium compounds (QACs) (Jia et al., 2022).

Mutations in these genes were responsible for low-level changes in antimicrobial susceptibility, however we found no evidence that exposure to each preservative caused significant cross- resistance to antibiotics. Characterisation of the SNPs found within *crp* and *rpoS* will allow a more accurate understanding of how these common mutations affect susceptibility to these preservatives, and whether the trade-off between biofilm formation and preservative adaptation can be exploited to reduce contamination in the food chain.

Biofilm populations continuously exposed to KCl or SL selected for mutations in both *rpoS* and *nlpD*. Deletion of either of these genes resulted in reduced growth in the presence of KCl relative to the wild type, but increased biofilm formation in the presence of SL. There was also reduced biofilm formation following deletion of either of these genes without preservative treatment and in the presence of NaCl. This indicates *rpoS* and *nlpD* affect susceptibility in a preservative- dependent manner, supporting our finding that these is no common mechanism of preservative tolerance. The *rpoS* and *nlpD* genes are located next to each other in an operon, which may indicate a shared regulator or function in response to each distinct preservative stress. The antagonistic action of these preservatives could be exploited as a preservative cocktail to growth and biofilm formation. Preservative combinations should be tested in our model to predict how this affects the evolutionary trajectory of planktonic and biofilm populations under these stresses.

The implications of this study support continued use of these preservatives in the food industry. Reduced biofilm formation and no evidence of cross-resistance to antibiotics following preservative exposure suggests adaptation when it does occur is unlikely to have negative impacts on food safety or section of AMR. This work can however be expanded in future to expand the range of selective conditions to better mimic industry-relevant biotic and abiotic surfaces and allow a more holistic interpretation of how preservatives affect biofilm formation of *S*. Typhimurium or other foodborne pathogens.

## Notes

**Funding** The authors gratefully acknowledge the support of the Biotechnology and Biological Sciences Research Council (BBSRC); JAA, ERH, ET and MAW were supported by the BBSRC Institute Strategic Programme Microbes in the Food Chain BB/R012504/1 and its constituent project BBS/E/F/000PR10349.

**Data access statement** The data that support the findings of this study are available on request from the corresponding authors.

### Competing Interest Statement

The authors have declared no competing interest.

