## Supplementary tables 1,2 for "Common food preservatives impose distinct selective pressures on *Salmonella* Typhimurium planktonic and biofilm populations"

**Supplementary Information**

**Supplementary table 1:** Minimum inhibitory concentrations (MICs, µg/mL) of different antibiotics in S. Typhimurium biofilm populations continuously exposed to sodium chloride (NaCl), potassium chloride (KCl), sodium lactate (SL) or sodium nitrite (SN) for 12 passages. The error margins of this test are one $log_2$-fold change from the wild type (WT). Results are representative of two biological and two technical replicates.

| **Populations continuously exposed to** | **NaCl** | | | | **KCl** | | | | **SL** | | | | **SN** | | | | **WT** |
|---|---|---|---|---|---|---|---|---|---|---|---|---|---|---|---|---|---|
| **Lineage** | **A** | **B** | **C** | **D** | **A** | **B** | **C** | **D** | **A** | **B** | **C** | **D** | **A** | **B** | **C** | **D** | |
| Chloramphenicol | 4 | 4 | 4 | 4 | 4 | 4 | 4 | 4 | 4 | 4 | 4 | 4 | 2 | 2 | 2 | 2 | 2 |
| Tetracycline | 0.5 | 0.5 | 0.5 | 0.5 | 0.5 | 0.5 | 0.5 | 0.5 | 1 | 1 | 1 | 1 | 1 | 1 | 1 | 1 | 0.5 |
| Azithromycin | 8 | 8 | 8 | 8 | 8 | 8 | 8 | | 8 | 8 | 8 | 8 | 8 | 8 | 8 | 8 | 4 |
| Ciprofloxacin | 0.008 | 0.008 | 0.008 | 0.008 | 0.008 | 0.008 | 0.008 | 0.008 | 0.016 | 0.016 | 0.125 | 0.016 | 0.008 | 0.008 | 0.008 | 0.008 | 0.016 |
| Kanamycin | 8 | 8 | 8 | 8 | 8 | 8 | 8 | 8 | 8 | 4 | 4 | 4 | 8 | 8 | 8 | 8 | 4 |
| Cefotaxime | 0.125 | 0.125 | 0.125 | 0.125 | 0.125 | 0.125 | 0.125 | 0.125 | 0.0625 | 0.0625 | 0.063 | 0.125 | 0.03125 | 0.03125 | 0.031 | 0.031 | 0.031 |
| Colistin | 0.5 | 1 | 1 | 0.5 | 0.5 | 1 | 0.5 | 0.5 | 0.5 | 0.5 | 0.5 | 0.5 | 1 | 1 | 1 | 1 | 1 |
| Ampicillin | 1 | 1 | 1 | 1 | 2 | 2 | 2 | 2 | 0.5 | 0.5 | 0.5 | 0.5 | 1 | 1 | 1 | 1 | 2 |

**Supplementary table 2:** Locations and descriptions of mutations found in planktonic and biofilm cultures exposed to NaCl, KCl, SL or SN, including unexposed biofilm controls. SNPs were identified by comparing FASTQ files from each isolate to *S.* Typhimurium 14028*S* reference genome CP001363 using Snippy version 4.6.

| **Drug** | **Lineage** | **Gene name** | **locus tag** | **Mutation** | **Function** |
|---|---|---|---|---|---|
| KCl | B | | STM14_0634 | missense_variant c.143C>A p.Pro48His | Unknown |
| NaCl | B | | STM14_1524 | synonymous_variant c.87G>A p.Thr29Thr | Unknown |
| NaCl | Control | | STM14_1524 | synonymous_variant c.87G>A p.Thr29Thr | Unknown |
| SN | C | | STM14_3103 | missense_variant c.1007C>T p.Ala336Val | Putative anaerobic dimethylsulfoxide reductase |
| SL | D | | STM14_4510 | missense_variant c.445T>A p.Trp149Arg | Unknown |
| SL | D | | STM14_4510 | missense_variant c.445T>A p.Trp149Arg | Unknown |
| SL | D | | STM14_4510 | missense_variant c.445T>A p.Trp149Arg | Unknown |
| NaCl | Planktonic | | STM14_0050 | missense_variant c.1691G>A p.Cys564Tyr | Putative glycosyl hydrolase |
| NaCl | Planktonic | | STM14_2943 | missense_variant c.152C>T p.Ala51Val | Unknown |
| KCl | B | amiC | STM14_3606 | missense_variant c.209G>T p.Arg70Leu | Murein separation cell division |
| KCl | B | amiC | STM14_3606 | missense_variant c.209G>T p.Arg70Leu | Murein separation cell division |
| KCl | B | amiC | STM14_3606 | missense_variant c.209G>T p.Arg70Leu | Murein separation cell division |
| NaCl | C | chbF | STM14_1598 | frameshift_variant c.847_848delGC p.Ala283fs | Chitobiose degradation |
| NaCl | C | chbF | STM14_1598 | frameshift_variant c.847_848delGC p.Ala283fs | Chitobiose degradation |
| SN | A | chbR | STM14_1597 | synonymous_variant c.762T>C p.Ser254Ser | Regulator of chitobiose degradation |
| SL | C | cpdA | STM14_3856 | frameshift_variant c.500dupC p.Leu168fs | cAMP phosphodiesterase |
| SL | C | cpdA | STM14_3856 | stop_gained c.659C>A p.Ser220* | cAMP phosphodiesterase |
| NaCl | A | crp | STM14_4173 | missense_variant c.56G>A p.Cys19Tyr | cAMP receptor protein |
| NaCl | A | crp | STM14_4173 | missense_variant c.56G>A p.Cys19Tyr | cAMP receptor protein |
| NaCl | A | crp | STM14_4173 | missense_variant c.56G>A p.Cys19Tyr | cAMP receptor protein |
| KCl | A | crp | STM14_4173 | missense_variant c.56G>A p.Cys19Tyr | cAMP receptor protein |
| KCl | A | crp | STM14_4173 | missense_variant c.56G>A p.Cys19Tyr | cAMP receptor protein |
| KCl | Planktonic | crp | STM14_4173 | missense_variant c.56G>A p.Cys19Tyr | cAMP receptor protein |
| SL | Planktonic | crp | STM14_4173 | missense_variant c.56G>A p.Cys19Tyr | cAMP receptor protein |
| SL | Planktonic | crp | STM14_4173 | missense_variant c.56G>A p.Cys19Tyr | cAMP receptor protein |

| | | | | | |
|---|---|---|---|---|---|
| SN | Planktonic | cspB | STM14_2420 | missense_variant c.40C>T p.Pro14Ser | Putative cold-shock protein |
| SL | C | dgcE | STM14_2620 | missense_variant c.1301C>A p.Ala434Glu | Cyclic d-GMP diguanylate cyclase |
| SN | B | dgcE | STM14_2620 | missense_variant c.2906G>A p.Gly969Asp | Cyclic d-GMP diguanylate cyclase |
| SL | B | ecnA | STM14_5215 | missense_variant c.70G>A p.Ala24Thr | Starvation adaptation TA system |
| SL | B | ecnA | STM14_5215 | missense_variant c.70G>A p.Ala24Thr | Starvation adaptation TA system |
| SL | B | ecnA | STM14_5215 | missense_variant c.70G>A p.Ala24Thr | Starvation adaptation TA system |
| SN | C | fimA | STM14_0635 | synonymous_variant c.441C>T p.Gly147Gly | Fimbriae |
| NaCl | C | flhC | STM14_2340 | frameshift_variant c.79_80insT p.Gln27fs | Flagella master regulator |
| NaCl | C | flhC | STM14_2340 | frameshift_variant c.79_80insT p.Gln27fs | Flagella master regulator |
| NaCl | Control | flhC | STM14_2340 | frameshift_variant c.79_80insT p.Gln27fs | Flagella master regulator |
| NaCl | Control | flhC | STM14_2340 | frameshift_variant c.79_80insT p.Gln27fs | Flagella master regulator |
| KCl | Control | flhC | STM14_2340 | frameshift_variant c.79_80insT p.Gln27fs | Flagella master regulator |
| KCl | Control | flhC | STM14_2340 | frameshift_variant c.79_80insT p.Gln27fs | Flagella master regulator |
| SN | C | flhC | STM14_2340 | missense_variant c.103C>T p.Leu35Phe | Flagella master regulator |
| SN | Control | flhD | STM14_2341 | frameshift_variant c.169delA p.Met57fs | Flagella master regulator |
| SN | D | flhD | STM14_2341 | stop_gained c.274C>T p.Gln92* | Flagella master regulator |
| SN | Planktonic | fliG | STM14_2391 | stop_gained c.628C>T p.Gln210* | Flagellar motor switch protein G |
| SN | B | galT | STM14_0900 | missense_variant c.448G>A p.Val150Ile | Galactose degradation |
| SN | D | gltB | STM14_4020 | missense_variant c.3704T>C p.Phe1235Ser | Glutamate synthase |
| NaCl | B | gltS | STM14_4511 | missense_variant c.644T>A p.Leu215Gln | Glutamate:sodium symporter |
| NaCl | Control | gltS | STM14_4511 | missense_variant c.644T>A p.Leu215Gln | Glutamate:sodium symporter |
| KCl | Control | gltS | STM14_4511 | missense_variant c.628C>T p.Arg210Cys | Glutamate:sodium symporter |
| SN | A | gltS | STM14_4511 | missense_variant c.584A>G p.Asp195Gly | Glutamate:sodium symporter |
| SN | A | gltS | STM14_4511 | missense_variant c.584A>G p.Asp195Gly | Glutamate:sodium symporter |
| NaCl | Planktonic | hnr | STM14_2119 | frameshift_variant c.860dupA p.Asn287fs | Response regulator of RpoS |
| SL | Planktonic | hupA | STM14_5011 | missense_variant c.47T>G p.Leu16Arg | Regulator HU subunit alpha |
| SN | D | idnR | STM14_5377 | missense_variant c.272G>A p.Gly91Asp | Regulator of gluconate metabolism |
| SN | D | lapC | STM14_2754 | missense_variant c.1345G>A p.Ala449Thr | LPS signal transducer |
| NaCl | A | lpxO | STM14_5156 | missense_variant c.773G>T p.Arg258Leu | LPS core subunit |
| NaCl | A | lpxO | STM14_5156 | missense_variant c.773G>T p.Arg258Leu | LPS core subunit |

| | | | | | |
|---|---|---|---|---|---|
| SL | A | malT | STM14_4234 | synonymous_variant c.1092G>C p.Ala364Ala | Regulator of maltose metabolism |
| NaCl | D | menA | STM14_4918 | missense_variant c.646A>C p.Thr216Pro | Menaquinone biosynthesis |
| KCl | B | nlpD | STM14_3527 | synonymous_variant c.630C>A p.Gly210Gly | Activates amiC during cell division |
| KCl | B | nlpD | STM14_3527 | synonymous_variant c.630C>A p.Gly210Gly | Activates amiC during cell division |
| KCl | B | nlpD | STM14_3527 | synonymous_variant c.630C>A p.Gly210Gly | Activates amiC during cell division |
| SL | B | nlpD | STM14_3527 | frameshift_variant c.598delG p.Ala200fs | Activates amiC during cell division |
| SL | B | nlpD | STM14_3527 | frameshift_variant c.598delG p.Ala200fs | Activates amiC during cell division |
| SL | B | nlpD | STM14_3527 | frameshift_variant c.598delG p.Ala200fs | Activates amiC during cell division |
| NaCl | Planktonic | ompF | STM14_1130 | missense_variant c.223G>A p.Gly75Ser | Outer membrane porin F |
| KCl | Planktonic | ompR | STM14_4217 | frameshift_variant c.129dupG p.Leu44fs | Osmolarity response regulator |
| NaCl | B | panF | STM14_4079 | missense_variant c.599T>A p.Leu200Gln | Pantothenate:Na+ symporter |
| NaCl | Control | panF | STM14_4079 | missense_variant c.599T>A p.Leu200Gln | Pantothenate:Na+ symporter |
| KCl | B | pphA | STM14_2241 | missense_variant c.101G>A p.Cys34Tyr | Misfolded protein stress response |
| KCl | B | pphA | STM14_2241 | missense_variant c.101G>A p.Cys34Tyr | Misfolded protein stress response |
| SN | D | prpR | STM14_0429 | synonymous_variant c.1068A>G p.Gly356Gly | Propionate catabolism operon regulatory protein |
| SN | Planktonic | ptsG | STM14_1377 | missense_variant c.529A>G p.Thr177Ala | Glucose-specific PTS system IIBC |
| NaCl | Planktonic | rfbK | STM14_2577 | missense_variant c.1171G>A p.Asp391Asn | Phosphomannomutase |
| SN | A | rlmJ | STM14_4326 | missense_variant c.227C>T p.Ala76Val | rRNA methyltransferase |
| NaCl | Planktonic | rna | STM14_0718 | frameshift_variant c.432delG p.Asn145fs | Ribonuclease I |
| SL | D | rpoD | STM14_3888 | missense_variant c.469C>G p.Leu157Val | RNA polymerase sigma factor |
| SL | D | rpoD | STM14_3888 | missense_variant c.469C>G p.Leu157Val | RNA polymerase sigma factor |
| SL | D | rpoD | STM14_3888 | missense_variant c.469C>G p.Leu157Val | RNA polymerase sigma factor |
| NaCl | C | rpoS | STM14_3526 | missense_variant c.812G>C p.Arg271Pro | Sigma factor, stringent response regulator |
| NaCl | D | rpoS | STM14_3526 | missense_variant c.812G>C p.Arg271Pro | Sigma factor, stringent response regulator |
| KCl | C | rpoS | STM14_3526 | missense_variant c.358A>C p.Ile120Leu | Sigma factor, stringent response regulator |
| KCl | C | rpoS | STM14_3526 | missense_variant c.358A>C p.Ile120Leu | Sigma factor, stringent response regulator |
| SL | A | rpoS | STM14_3526 | stop_gained c.347T>A p.Leu116* | Sigma factor, stringent response regulator |
| SL | A | rpoS | STM14_3526 | missense_variant c.875G>A p.Gly292Asp | Sigma factor, stringent response regulator |
| SL | A | rpoS | STM14_3526 | stop_gained c.347T>A p.Leu116* | Sigma factor, stringent response regulator |
| SL | C | rpoS | STM14_3526 | frameshift_variant c.921delT p.Glu308fs | Sigma factor, stringent response regulator |

| | | | | | |
|---|---|---|---|---|---|
| SL | Planktonic | rpoS | STM14_3526 | conservative_inframe_deletion c.598_603delGAGCAA p.Glu200_Gln201del | Sigma factor, stringent response regulator |
| SL | Planktonic | rpoS | STM14_3526 | conservative_inframe_deletion c.598_603delGAGCAA p.Glu200_Gln201del | Sigma factor, stringent response regulator |
| KCl | C | rpoZ | STM14_4506 | frameshift_variant c.99_100dupCG p.Gly34fs | RNA polymerase subunit |
| KCl | A | rpoZ | STM14_4506 | synonymous_variant c.30A>C p.Val10Val | RNA polymerase subunit |
| KCl | C | rpoZ | STM14_4506 | frameshift_variant c.37delA p.Ile13fs | RNA polymerase subunit |
| KCl | C | rpoZ | STM14_4506 | frameshift_variant c.37delA p.Ile13fs | RNA polymerase subunit |
| NaCl | D | rseA | STM14_3233 | missense_variant c.301C>T p.Leu101Phe | Anti-rpoE factor |
| SN | B | scsB | STM14_1267 | missense_variant c.1459T>C p.Ser487Pro | Suppression of copper sensitivity protein |
| SN | D | sipB | STM14_3484 | missense_variant c.1031C>T p.Ala344Val | Translocation machinery component |
| KCl | D | stcD | STM14_2652 | missense_variant c.155A>C p.Asp52Ala | Unknown |
| KCl | D | stcD | STM14_2652 | missense_variant c.155A>C p.Asp52Ala | Unknown |
| KCl | D | stcD | STM14_2652 | missense_variant c.395C>T p.Thr132Ile | Unknown |
| SL | Planktonic | tatB | STM14_4780 | stop_gained c.178C>T p.Gln60* | Sec-independent translocase |
| SN | C | yacH | STM14_0189 | missense_variant c.691A>G p.Thr231Ala | Unknown |
| NaCl | Planktonic | ychP | STM14_2138 | missense_variant c.742C>T p.His248Tyr | Unknown |
| SN | B | yciC | STM14_2097 | missense_variant c.266G>A p.Gly89Glu | Unknown |
| SN | B | yciC | STM14_2097 | missense_variant c.266G>A p.Gly89Glu | Unknown |
| KCl | B | ycjF | STM14_2034 | synonymous_variant c.564C>T p.Val188Val | Unknown |
| KCl | B | ycjF | STM14_2034 | synonymous_variant c.564C>T p.Val188Val | Unknown |
| KCl | B | ycjF | STM14_2034 | synonymous_variant c.564C>T p.Val188Val | Unknown |
| SN | B | yehY | STM14_2668 | missense_variant c.928C>T p.Pro310Ser | ABC transporter subunit |
| SN | C | yhbW | STM14_3958 | missense_variant c.73G>A p.Ala25Thr | Monooxygenase |
| SN | C | yhcC | STM14_4017 | synonymous_variant c.630T>C p.Gly210Gly | Oxidoreductase |
| KCl | D | yjdB | STM14_5165 | frameshift_variant c.743delG p.Gly248fs | Unknown |
| KCl | D | yjdB | STM14_5165 | frameshift_variant c.743delG p.Gly248fs | Unknown |
| KCl | D | yjdB | STM14_5165 | frameshift_variant c.743delG p.Gly248fs | Unknown |
| NaCl | D | yjfN | STM14_5260 | missense_variant c.208C>G p.Arg70Gly | Activates degP proteolysis |
